# Anatomical and functional analysis of the corticospinal tract in an FRDA mouse model

**DOI:** 10.1101/2024.06.28.601178

**Authors:** Misa Nishiyama, John Kalambogias, Fumiyasu Imai, Emily Yang, Sonia Lang, Joriene C de Nooij, Yutaka Yoshida

**Affiliations:** Burke Neurological Institute, White Plains, New York, United States; Department of Neurology, Columbia University, New York, NY, USA; Brain and Mind Research Institute, Weill Cornell Medicine, New York, United States; Neural Circuit Unit, Okinawa Institute of Science and Technology Graduate University, Okinawa, Japan

## Abstract

Friedreich’s ataxia (FRDA) is one of the most common hereditary ataxias. It is caused by a GAA repeat in the first intron of the FXN gene, which encodes an essential mitochondrial protein. Patients suffer from progressive motor dysfunction due to the degeneration of mechanoreceptive and proprioceptive neurons in dorsal root ganglia (DRG) and cerebellar dentate nucleus neurons, especially at early disease stages. Postmortem analyses of FRDA patients also indicate pathological changes in motor cortex including in the projection neurons that give rise to the cortical spinal tract (CST). Yet, it remains poorly understood how early in the disease cortical spinal neurons (CSNs) show these alterations, or whether CSN/CST pathology resembles the abnormalities observed in other tissues affected by FXN loss. To address these questions, we examined CSN driven motor behaviors and pathology in the YG8JR FRDA mouse model. We find that FRDA mice show impaired motor skills, exhibit significant reductions in CSN functional output, and, among other pathological changes, show abnormal mitochondrial distributions in CSN neurons and CST axonal tracts. Moreover, some of these alterations were observed as early as two months of age, suggesting that CSN/CST pathology may be an earlier event in FRDA disease than previously appreciated. These studies warrant a detailed mechanistic understanding of how FXN loss impacts CSN health and functionality.

## Introduction

Friedreich’s ataxia (FRDA) is a multisystem disorder, characterized by progressive ataxia, scoliosis, loss of sensation, increased incidence of diabetes, and hypertrophic cardiomyopathy (Koeppen and Mazurkiewicz, 2013; Rojas et al., 2021; Keita et al., 2022; Lees et al., 2022). It is caused by an abnormal elongation of the GAA repeat in the first intron in the FXN gene, which encodes Frataxin, a small mitochondrial protein (Campuzano et al., 1996). On average, patients start experiencing motor deficits between 11-15 years of age and become unable to care for themselves some 15-20 years after disease onset. In general, however, the cause of death is heart failure associated with cardiac muscle hypertrophy (Koeppen, 2011; Tsou et al., 2011).

Affected individuals usually carry two alleles with GAA repeat expansions – sometimes reaching as many as 1700 repeats (Campuzano et al., 1996; Dürr et al., 1996). For comparison, unaffected individuals typically have 9 -35 repeats in the FXN gene. Previous studies demonstrated that a larger number of GAA repeats correlates with earlier disease onset and increased severity of symptoms, indicating that a patient’s disease course is largely dictated by the allele with the smallest number of repeats (Dürr et al., 1996; Delatycki and Bidichandani, 2019). The GAA repeat expansion causes epigenetic alterations that interfere with normal FXN expression (Herman et al., 2006; Yandim et al., 2013; Chutake et al., 2015; Rodden et al., 2022). Although different tissues appear to express different levels of FXN, on average FXN protein levels in patients are reduced to 5-35% of the levels in unaffected individuals (Campuzano et al., 1997; Willis et al., 2008; Deutsch et al., 2010; Saccà et al., 2011; Guo et al., 2018).

Frataxin (Fxn) is implicated in several mitochondrial and cellular functions (Rötig et al., 1997; Culley et al., 2023). Within mitochondria, it is important in the formation of Fe-S clusters which, in turn, serve as essential cofactors in multiple mitochondrial (and cellular) enzymes, including Complex I, II, and III of the respiratory chain, and Aconitase of the TCA cycle (Read et al., 2021). Consequently, Fxn protein deficiencies result in mitochondrial dysfunction, defects in iron homeostasis, and increased oxidative stress (Chen et al., 2002; Mühlenhoff et al., 2002). Despite the requirement of mitochondrial function in all tissues, a loss in Fxn most prominently affects tissues with high metabolic needs, including heart muscle and certain neural tissues. Indeed, the loss of proprioceptive deep tendon reflexes is an early and signature event in the disease process(Geoffroy et al., 1976). Another neural tissue that is impacted early in the disease is the dentate nucleus in cerebellum (Koeppen et al., 2007; Lin et al., 2017).

More recently, FRDA-associated motor deficits due to cortical pathology, in particular of cortical spinal neurons (CSNs) have garnered increased attention. CSNs are located in layer Vb in the cerebral cortex and project to inter- and motor neurons in the spinal cord through the cortical spinal tract (CST) (Campuzano et al., 1996; Koeppen et al., 2009, 2011; Koeppen, 2011). The CSN/CST is a major pathway involved in voluntary movements and is especially important for fine or skilled motor control such as required for writing or typing (Lawrence and Kuypers, 1968; Sterr et al., 2014). Lesions related to the CSNs/CST that have been reported following post-mortem analyses in FRDA patients include thinning of the motor cortex, atrophy and loss of Betz cells, as well as CST atrophy and demyelination in the spinal cord (Koeppen and Mazurkiewicz 2013; Koeppen et al. 2017; Lamarche, Lemieux, and Lieu 1984; Rezende et al. 2016; Harding et al. 2021). Analysis of the cerebral cortex in mice, not limited to CSNs, also reported increased iron accumulation and oxidative stress following a localized deletion of Fxn (Chen et al. 2016). In addition, using a finger tapping (FT) test designed to assess CST function, FRDA patients showed significant decreases in FT velocity without loss of regularity or decrease in amplitude (Naeije et al. 2021).Abnormalities in CSN/CST function are thought to be a late FRDA disease phenotype and are difficult to entangle from other motor deficits. As such, studies using animal models have mainly focused on the cerebellum and dorsal root ganglia (DRG). Yet, with increasing numbers of therapeutic interventions being evaluated in preclinical and clinical trials (including gene therapy and supplementation of Fxn via viral vectors), it is critical to consider the consequences for all FRDA disease phenotypes. Therefore, a more detailed understanding of the temporal and mechanistic aspects of CSN/CST dysfunction in FRDA is urgently needed.

We here used a variant of the recently developed YG8JR FRDA mouse model to examine the consequences of the loss of Fxn on CSNs and the CST. Previous studies demonstrated that this mouse model exhibits impaired motor function from an early age, resembling what is observed in FRDA patients (Gérard et al., 2023; Kalef-Ezra et al., 2023). Consistent with these observations, our morphological, functional, and behavioral analysis demonstrate that CSNs/CST functional output is significantly impaired in YG8JRex4 mice, even by 8-12 weeks of age. Thus, together our studies offer critical new insights into CSN/CST dysfunction and pathology in FRDA. They also indicate that CSN/CST impairments may contribute earlier to the FRDA-associated motor phenotype than previously recognized.

## Materials and methods

### Animals and animal husbandry

*Fxn*^em2.1Lutzy^ Tg(FXN)YG8Pook/800J mice, generally designated as YG8JR, were obtained from The Jackson Laboratory (stock number #030395)(Gérard et al., 2023; Kalef-Ezra et al., 2023). They were crossed with *Fxn*^Δ4/+^ mice which were obtained from Dr. Jordi Magrane (Weill Cornell Medicine) to replace the exon 2 deletion in YG8JR animals with the exon 4 mutant allele (Cossée et al., 2000). After establishing the line, the mouse colony was maintained by breeding *Fxn*^Δ4/+^; Tg(FXN)YG8Pook/800J males and females. *Fxn*^Δ4/Δ4^; Tg(FXN)YG8Pook/800J mice were used as FRDA animals; *Fxn*^+/+^ , *Fxn*^Δ4/+^ , *Fxn*^Δ4/+^;Tg(FXN)YG8Pook/800J, or *Fxn*^Δ4/+^ ;Tg(FXN)YG8Pook/800J were all used as littermate control mice. FRDA and control mice were identified using tail DNA genotyping using a common forward primer for exon4: Exon4-common forward: 5’ CTG TTT ACC ATG GCT GAG ATC TC, and two reverse primers specific for either the wild-type allele, Exon4-wild type reverse: 5’ CCA AGG ATA TAA CAG ACA CCA TT, or the exon4 knockout allele, exon4-knockout reverse: 5’CGC CTC CCC TAC CCG GTA GAA TTC (Miranda et al., 2002). For the YG8 transgene, the following primers were used: YG8 transgene forward 5’ GCAAAAGCTGGAGATCAAAGTGTGA, and YG8 transgene reverse 5’ GAGGCAACACATTCTTTCTACAGA (Sarsero et al., 2004). Animals with the YG8 transgene were also routinely assessed for GAA repeat length as described below (see Fig. 1B). Mice were maintained on a 12 hr light/dark cycle, with food and water ad libitum, except during training periods for behavioral experiments (see below). All animals were treated according to institutional and National Institutes of Health guidelines, with the approval of the Institutional Animal Care and Use committee at Burke Neurological Institute/Weill Cornell Medicine.

**Fig. 1.**
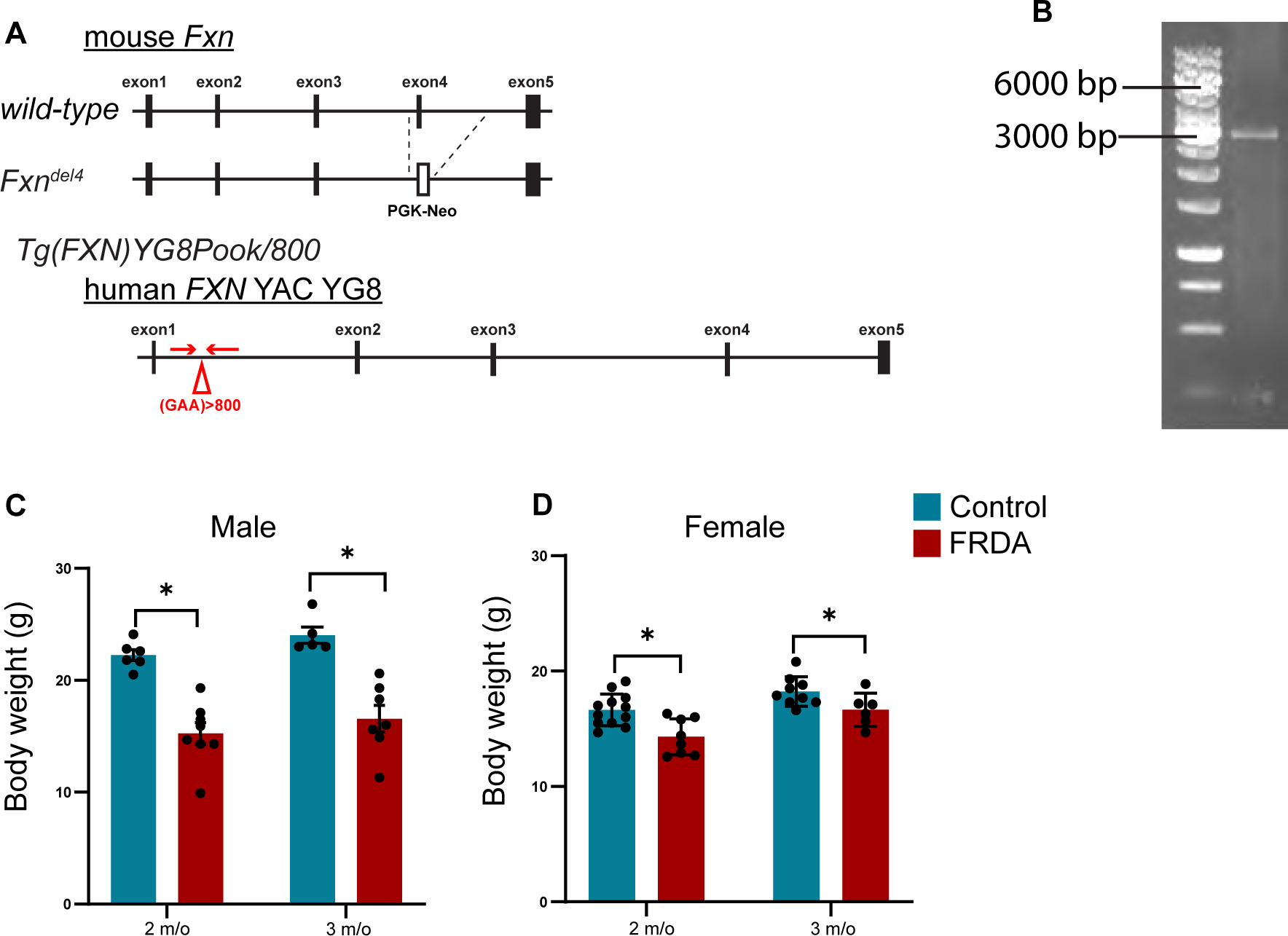
Body weight of FRDA mice is reduced. (A), Allele structure of mouse FRDA^Δ4^ and Tg(FXN)YG8Pook/800. FRDA^Δ4^ allele lacks exon4. T Tg(FXN)YG8Pook/800 contains the human *FXN* gene from YAC YG8. More than 800 repeats of GAA are inserted into intron1. Arrows indicates primers for GAA repats. (B), PCR analysis of GAA repeats in Tg(FXN)YG8Pook/800. The band around 3000 bp corresponds to 800 repeats. (Fig.1B).(C, D), Body weight of 2 and 3 months old male (C) and female (D) mice of (n= 5-7, for males, n=6-12 for females, Mean ± SD). *p<0.05 (Student’s t test-test).

### GAA repeat analysis

DNA was extracted from tails (digested with proteinase K buffer) with standard phenol-chloroform extraction and ethanol precipitation. PCR amplification was carried out using Premix D mixture of the FailSafe™ PCR System (Epicentre, # FS99250). The following primers were used for this analysis: FRDA-104F forward 5’ GGCTTGAACTTCCCACACGTGTT, and FRDA-629R Reverse 5’ AGGACCATCATGGCCACACTT (Bidichandani, Ashizawa, and Patel 1998). PCR products were run on 1% agarose gels for one hour. A band migrating around 3000 bp corresponds to 800 repeats (Fig.1B).

### Motor behavioral testing

Behavioral tests were performed with two and/or three month old mice. For each test, mice were habituated to the environment and trained on the test prior to recording the data. For each test, training was performed equally across all mice. Except when indicated otherwise, tests were performed a minimum of four times per animal and the recorded data was the average of all trials. Individual tests are described below. Some tests were only performed with male mice given that FRDA females were often too small and fragile to tolerate the test or the required food deprivation.

#### Open field test

Male mice were placed in a 40 cm x 40 cm chamber and the total locomotor distance within 10 min was measured with Any-maze tracking software (ANY-maze, Stoelting Co., USA).

*Grip strength.* Forelimb grip strength was measured by a Grip Strength Meter (Columbus Instruments, GRIP STRENGTH METER, USA). Maximum strength was measured and averaged across four trials.

#### Rotarod test

Rotarod test was carried out with an Ugo basile accelerating rotarod (Ugo basile, Italy). Four trials were performed with the speed of the rotation gradually increasing from 4 to 40 rpm in 400 second with 3 min intervals. The latency to fall from the rod was recorded.

#### Treadmill test

Treadmill test was carried out with an accelerating treadmill (Harvard Apparatus). To stimulate continued locomotion, mice received a mild electric shock (0.3 mA) if they reach the rear end of the treadmill. After 3 min of warm-up at a rate of 15 cm/s, treadmill speed was increased from 15 to 80 cm/second, in 5 cm/second increments. The mice were allowed to run for 20 second at each speed. When the mice received 3 electric shocks, the prior speed was recorded as the maximum speed. Maximum speed was measured in over four trials with 3 min intervals between trials.

#### Beam walking test

The beam walking test was carried out with 70 cm long round wooden poles, either 25.4 mm or 12.7 mm in diameter. After two training runs, mice were placed on one end of the pole and the time to cross to the other side, as well as the number of miss steps, was recorded. Each trial was conducted four times per pole diameter. Trials were video recorded by cameras (Sony 4K FDR-AX43) for analysis.

#### Horizontal ladder test

The horizontal ladder test was carried out with a 70 cm long ladder. For regular ladder walking, rungs were spaced 2 cm apart. For irregular ladder walking, rungs were randomly spaced 1 to 3 cm apart. (The pattern was fixed throughout the experiment). Mice were trained for two consecutive days by allowing them to cross the ladder twice on each training day. The test was conducted over two days, with mice allowed to walk on the ladder a maximum of five times per test day. Two trials with consistent pacing (without stopping midway) were selected for analysis each day, resulting in a total of four trials used for analyses. The tests were video recorded by cameras (Sony 4K FDR-AX43) and time to cross the ladder and the number of missteps were measured.

#### Pellet reaching test

A single-pellet reaching task was carried out as previously reported with some modifications (Xu et al., 2009). Three month old male mice were kept under food restriction and body weight was monitored so not to drop below 80% of the original weight. During the first two days of habituation, mice were placed into a custom-made chamber with 20 sugar pellets for 20 min. In the training period, mice were allowed to reach and grasp a sugar pellet from a central opening within the holding cage. When mice were able to complete >20 trials within 20 min with at least 70% of trials performed by the same forepaw (to assess paw dominance), the mice were regarded as trained for the test. During the test period (7 days), sugar pellets were placed near the left or right opening (depending on the dominant forelimb of the mice) and the number of successful trials over 30 trials or within 20 min was recorded. Success rates of days 6 and 7 were averaged for comparative analysis across animals.

#### Pasta handling test

The pasta handling test was carried out as previously reported with some modifications (Ruder et al., 2021). Three month old male mice were kept under food restriction and body weight was monitored so not to drop below 80% of the original weight. During the habituation/training period, mice were placed into an 8 cm x 7 cm chamber and given four (2 cm) pasta sticks and allowed to eat for 20 min. During the test period, one pasta stick was given for three consecutive days. For each day, the time to complete eating the stick (maximum 20 min) and the number of times the stick was dropped were measured.

### Corticospinal neuron and mitochondrial labeling

Corticospinal neurons in motor cortex were retrogradely labeled with adeno associate virus (AAV) expressing a tdTomato reporter. Six to seven week old mice were anesthetized with isoflurane and placed on a stereotaxic frame. AAVretro-CAG-tdTomato (3.6 x 10^13^ GC/mL; Addgene, 0.3 µL/site) was injected into spinal cord level C6 (0.5 mm lateral from the midline, 0.5 and 1.0 mm depth from the spinal surface, 0.3 µL/site) using glass capillaries. For FRDA mice, spinal C6 injection sites were adjusted to accommodate their reduced size (0.45 mm lateral, 0.45 and 0.9 mm depth, 0.3 µL/site). To selectively label mitochondria in corticospinal neurons, AAVretro-CAG-Cre (4.3 x 10^12^ CG/mL; UNC vector core, 0.3 µL/site) was injected into the spinal cord at the C6 level (0.5 mm lateral, 0.5 and 1.0 mm depth for control; 0.45 mm lateral, 0.45 and 0.9 mm depth for FRDA). The following week, a 1:1 mixture of AAV8-hSyn-DIO-EGFP (2.3 x 10^13^ CG/mL, Addgene) and AAV8-Flex-3mts-mScarlet (1.14 x 10^13^ CG/mL) (Chertkova et al. 2017) was injected (0.3 µL/site) into the cortical hemisphere (AP 0 mm, ML 1.5 mm, DP 0.6 mm, AP 1.0mm, ML 1.5 mm, DP 0.5mm from bregma for control animals; AP 0 mm, ML 1.45 mm, DP0.45 mm, AP 0.9 mm, ML 1.45 mm, DP 0.45 mm for FRDA mice). Two weeks after AAV injections, mice were perfused with 4% PFA/PB and processed for histological analyses.

### Histological and immunological analyses

Mice were deeply anesthetized with ketamine and xylazine and then fixed by transcardial perfusion with PBS (10 ml) and 4% paraformaldehyde (PFA)/Phosphate buffer (PB) (20 ml). Brain and spinal cord were dissected free, post-fixed in 4% PFA/PB for overnight at 4°C, and (after washing in PBS) cryoprotected in 30% sucrose/PBS for 3 days. Muscles were post-fixed in 4% PFA/PB for 20 min at 4°C, and then placed in 30% sucrose/PBS solution for overnight. Tissues were embedded in Tissue-Tek OCT compound (Sakura Finetek) and sectioned at 50 μm (brain, spinal cord) or 16μm (muscle) using a Cryostat (Leica).

Sections were blocked in 1% BSA in PBST (0.1% tween20/PBS) for 30 min prior to antibody staining. Primary antibodies used were Cux1 (1:500, 11733-1-AP, Proteintec), CTIP2 (1:200, ab18465, ABCAM), NeuN (1:500, MAB377, Millipore), PKC gamma (1:500, ab71558, ABCAM), ChAT (1:200, AB144P, Millipore), vGluT1 (1:1000, AB5905, Millipore), Neurofilament (1:1000, AB1987, Millipore), and Synaptophysin (1:1000, 101104, Synaptic Systems). Primary Antibodies were diluted in PBST and incubated at room temperature for overnight. The following day, sections were rinsed with PBST and incubated in Alexa Fluor 488-, Cy3-, or Cy5-labeled secondary antibodies (Jackson ImmunoResearch) for two hours. To label neuromuscular junctions, alpha-Bungarotoxin Conjugate (1:1000, B1196, ThermoFisher Scientific) was added to the secondary antibody solution. Sections were washed with PBST three times and nuclei were labeled by DAPI (1:1000, D3571, Thermofisher scientific). Sections were mounted on glass slides with Vectashield (Vector Laboratories). Images were acquired using Leica (SP8) or Nikon (A1 HD25 / A1R HD25) confocal microscopes. For mitochondrial analysis, images were acquired with a Nikon A1 HD25 / A1R HD25 confocal microscope with a 60X/oil objective lens. Images were deconvoluted and denoised with Nikon software. Mitochondria in soma were trimmed with Imaris (Oxford Instruments). Total volume of mitochondria in each cell was analyzed with Mitochondria analyzer plugin for imageJ/Fiji (Chaudhry, Shi, and Luciani 2020). Block size and C-value were set as 3.5 and 10, respectively.

### EMG recordings

We recorded electromyographic (EMG) responses in the flexor carpi radialis (FCR) and bicep muscles bilaterally using percutaneous Nickel-chrome alloy wires (California Fine Wire [100189]) in response to motor cortex stimulation as previously reported (Gu, Serradj, et al. 2017; Gu, Kalambogias, et al. 2017).Anesthesia was induced using a ketamine/xylazine cocktail and were subsequently administered with ketamine to continue anesthesia and maintain appropriate muscle tone.

A craniotomy was conducted over the forelimb area of M1 where electrode penetrations were made perpendicular to the pial surface. We used Elgiloy microelectrodes (Microprobes; 0.1 Megohms impedance, 0.229 mm shaft diameter, 2-3 mm tip diameter). Trains of three pulse stimuli (0.2ms biphasic pulse, 33ms interstimulus interval for trains, every 2s, n=30 sweeps) were delivered to the contralateral motor cortex using a constant current stimulator. For stimulus triggered averaging, we used single pulse stimuli (0.2ms biphasic; every 0.5s, n=200 sweeps). EMG recordings were conducted employing a differential AC amplifier with low- and high-pass filtration (model 1700; AM Systems). Utilizing two wire electrodes within each muscle and a separate ground, we conducted differential recordings to confirm the adequacy of the EMG recordings and muscle placements, observing increased EMG activity with passive joint rotation. Subsequently, EMG signals were acquired through an analog-to-digital converter (CED) and processed utilizing Signal software (version 6.05; CED).

The threshold for three and single pulse stimulation was determined when nearly half of the stimulus trains were able to evoke muscle EMG responses. For three pulses, currents of 1, 1.1, 1.2 and 1.3 times the threshold were delivered every 2 s to generate recruitment curves. Ensemble averages of EMG responses were constructed from individual sites across each animal where recruitment curves were then calculated as the area under the multi-unit response curve (mV*ms) for each current value used. For stimulus triggered averaging, post-stimulus facilitation was generated by averaging EMG responses across 200 stimuli. Latency onset measurements were then determined following the first change of average EMG activity over the baseline.

### Statistical Analysis

Statistical evaluation was performed using Student’s t test, and values are shown as mean ± SD. For EMG analysis, statistical evaluation was performed using Student’s t test and Two-way ANOVA. p < 0.05 was considered significant.

## Results

### FRDA-YG8JR mice are smaller and develop lower gastrointestinal tract impairments

FRDA patients develop severe motor deficits but the extent to which these primarily result from proprioceptive and cerebellar dysfunction or may be caused by other central nervous system impairments, remains poorly understood. To study the contribution of cortical spinal motor neurons (CSNs) and the associated cortical spinal tract (CST) in FRDA-associated motor impairments, we obtained the *Fxn^em2·1Lutzy^ Tg(FXN)YG8Pook/800J*, mouse model, also referred to as YG8JR mice, from Jackson labs. Originally developed by the Pook lab, the YG8JR mice carry a null mutation for mouse Frataxin (a deletion of exon 2) along with a YAC transgene that comprises the human *FXN* gene with a GAA repeat expansion in intron 1 (Fig. 1A). In this FRDA mouse model, animals solely rely on the limited level of FXN protein expressed from the human allele. The original YG8sR line contained approximately 200-300 GAA repeats (Anjomani Virmouni et al., 2015), but has since been bred by Jackson labs to contain over 800 GAA repeats (Fig 1B). Recent studies have demonstrated that the levels of FXN protein are significantly reduced in these animals (Gérard et al., 2023; Kalef-Ezra et al., 2023) and that this animal model best reflects the course of disease observed in FRDA patients. Given an initial concern (later determined unfounded) over the exon 2 null mutation, we replaced the exon 2 allele with the exon 4 mutant allele (Cossée et al., 2000). To distinguish this mouse model from the YG8JR Jackson lab animals we termed them YG8JRex4, but for simplicity we will generally refer to them as “FRDA”.

As reported previously, we find that both male and female YG8JRex4 FRDA mice are significantly smaller than control mice by two and three months of age (Fig. 1C-D). In addition, we found that many FRDA mice (both male and female) developed rectal prolapse when they were around two months old. Indeed, by seven months of age, 70% of the male mice had developed rectal prolapse. Mice that developed rectal prolapse were euthanized in accordance with IACUC guidelines. Due to the high mortality (euthanasia rate) of older FRDA mice, we limited most of our analyses to two and three month old animals.

### FRDA mice exhibit impairments in skilled motor control

The CST is a major descending pathway from CSNs in primary motor cortex. The CSN cell bodies are located in layer 5b and project to the spinal cord, where, in humans, they form connections with interneurons and motor neurons. In rodents, direct connections to motor neurons are initially formed but are eliminated in adulthood (Yang and Lemon, 2003; Alstermark and Ogawa, 2004; Gu et al., 2017a). The CST is especially relevant during fine motor coordination(Lawrence and Kuypers, 1968; Duque et al., 2003; Sterr et al., 2014; Wang et al., 2017) . To begin to examine the extent of CSN/CST impairments in YG8JRex4 FRDA mice, we performed several behavioral assays testing both general and skilled motor behaviors. First, we assessed locomotor activity using the open field test. We found that during a 10 minute observation period, the total locomotor distance of FRDA mice was significantly decreased when compared to control mice (Fig. 2A). In addition, when assessing treadmill running, maximum running speed was significantly reduced in FRDA mice when compared to controls (Fig. 2B). We also measured grip strength as a means to assess forelimb strength. FRDA mice showed significantly weaker peak force compared to control mice (Fig. 2C). In contrast, FRDA mice performed similar to controls in the Rotarod test, suggesting they do not suffer from impaired motor coordination (Fig. 2D).

**Fig. 2.**
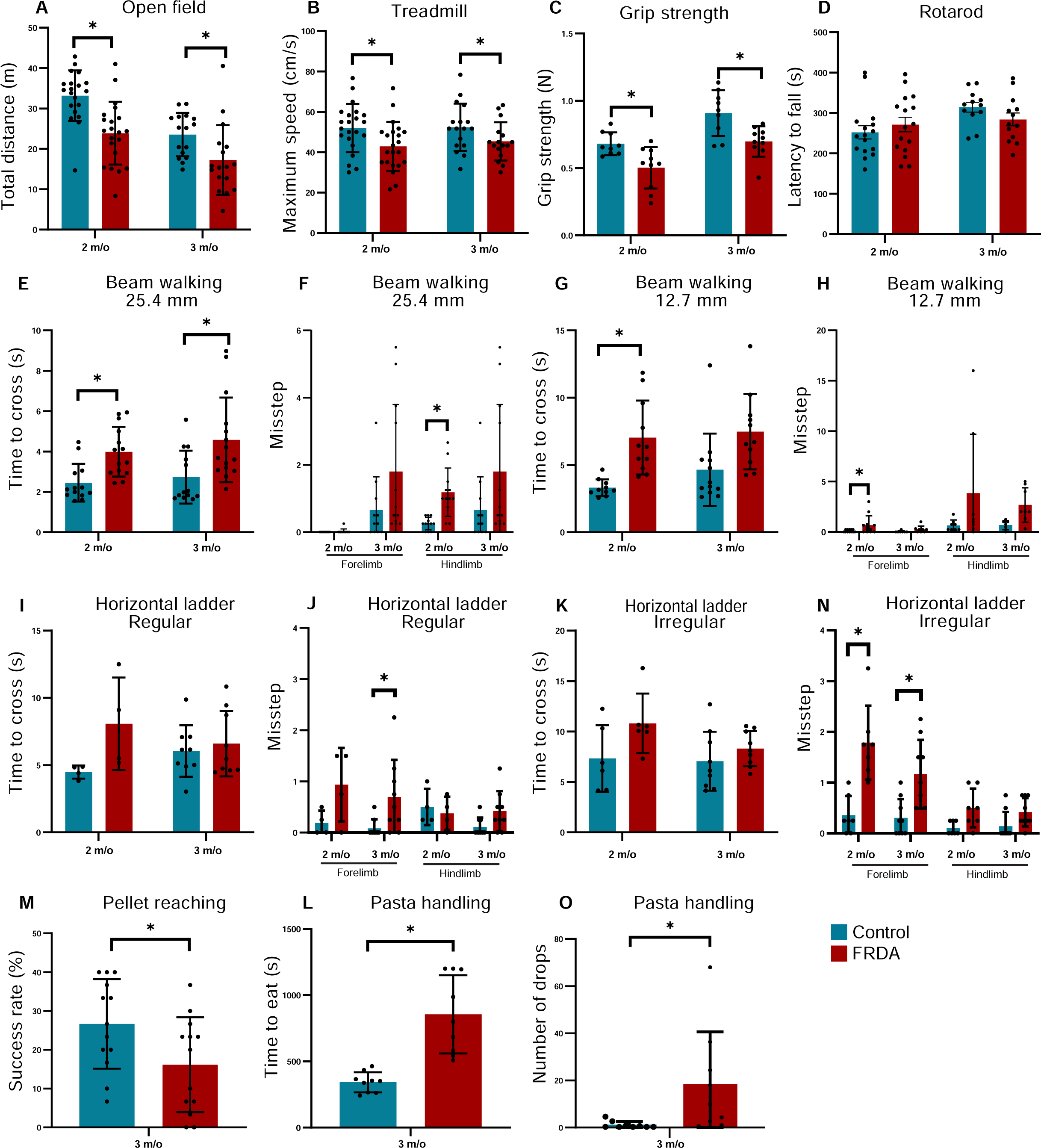
FRDA mice show motor deficit. (A), Traveled distance during open field test (n=16-21). (B), Maximum speed on the treadmill (n=16-22). (C), Grip strength of forelimb (n=9-10). (D), Latency to fall in accelerating Rotarod test (n=12-16). (E-H), Time (E, G) and missteps (F, H) to cross 25.4 mm (E, F, n=13-15) and 12.7 mm (G, H, n=10-12) diameter beams. (I-N), Time (I, K) and missteps (J, N) to cross regular (I, J, n=4-9) and Irregular (n=6-9) ladders. (M), Success rate of pellet reaching (n=13). (L-O), Time to eat a 2-cm pasta stick (L) and the number of pasta drops of the mice (O) during pasta handling test (n=9). All graphs are represented as Mean ± SD. *p<0.05 (Student’s t test).

To assess more specific motor behaviors, we performed the beam walking test (requiring precise foot placement and balance), pellet reaching tests (precision reach and digit control), and the pasta handling test (digit control). The beam walking test was conducted using beams of two different widths, 25.4 and 12.7 mm, respectively. Two and three month old FRDA mice both showed significantly longer time to cross the 25.4mm diameter pole compared to controls and also showed more missteps (Fig.2E-H). Similar results were observed for the 12.7 mm beam, but the difference only reached significance for the two month old animals. The pellet reaching test and pasta handling test were both conducted at three months of age, given that at two-months the FRDA mice seemed too frail for food restriction (part of the testing procedure). We find that FRDA mice showed lower success rates in the pellet reaching test compared to controls (Fig.2M). Similarly, FRDA mice needed considerably more time to eat the 2 cm stick of pasta and more frequently dropped the pasta on the floor when compared to control animals (Fig2L-O). Together these results indicate that FRDA mice exhibit general locomotor dysfunction but are particularly impaired in skilled motor behaviors.

### FRDA mice show significantly diminished CST output

To determine whether impairments in motor skill of FRDA mice could be a consequence of CS circuit dysfunction, we tested the functionality of connections between motor cortex and muscle. In mice, at forelimb levels, the CST indirectly activates motor neurons via interneurons in the cervical spinal cord, which in turn can elicit contraction of the muscles they innervate. Thus, to test CST output, we recorded motor evoked potentials (MEPs) in bicep and flexor carpi radialis (FCR) muscles in response to primary motor cortex (M1) stimulation (Fig. 3A). We first used stimulus-triggered averaging (Gu et al., 2017a), a single pulse microstimulation approach, to evaluate the timing of EMG responses in both bicep and FCR following M1 stimulation in both control and FRDA mice (Fig. 3B). At two months of age, for both biceps and FCR, we were unable to detect differences in latency to EMG onset between control and FRDA mice (Fig. 3C, D). Consistent with a disynaptic response and with previously reported latencies (Gu et al., 2017a), we find that EMG responses are observed after ∼13ms.

**Fig. 3.**
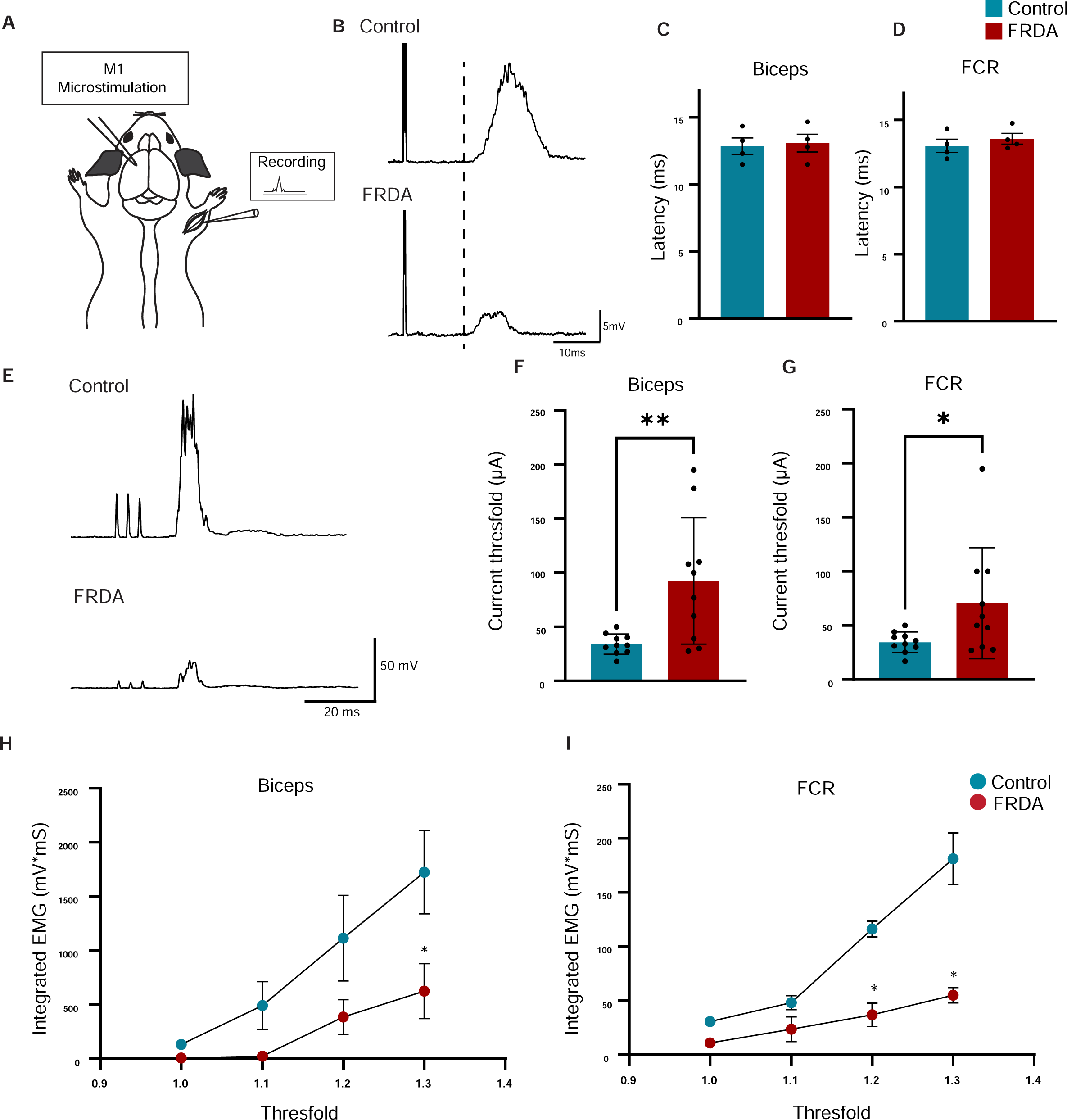
Motor cortex stimulation-evoked muscle responses display functional differences in FRDA mice. (A), Schematic drawing of EMG recordings from contralateral flexor carpi radialis (FCR) and biceps brachii muscles with primary motor cortex (M1) stimulation (single pulse for B-D and 3 pulses, every 2s for E-I) in 2 months old mice. (B), Representative EMG traces from FCR muscles of control (up) and FRDA (down) mice, after a single pulse (80mA). (C-D), Single pulse microstimulation latencies for biceps (C) and FCR (D) after M1 stimulation in control (n=4) and FRDA (n=4) mice. (E), Representative EMG traces from FCR muscles of control (up) and FRDA (down) mice after three stimulation pulses (32.2mA) (F, G), Three pulse microstimulation current thresholds for biceps (F) and FCR (G) following M1 stimulation in control (n=5, 2 sites) and FRDA (n=5, 2 sites) mice. (H-I). EMG recruitment curves from control and FRDA mice between 1x, 1.1x and 1.3x threshold for bicep (H) and FCR (I) (n=4 for control, n=5 for FRDA mice). ** p < 0.01, *p<0.05 (paired test, F, G). * p < 0.05 (two-way ANOVA with Bonferroni’s post hoc tests, H, I).

To further assay for muscle EMG responses and evaluate for CST functionality, we then used trains of three pulses to produce MEPs in controls and FRDA mice (Fig. 3E). We tested the threshold responses for M1 three pulse stimulation, which was defined when nearly half of the stimulation trains evoked a muscle EMG response (Fig. 3F,G). Threshold currents to evoke bicep and FCR EMG muscle responses following M1 stimulation were significantly increased (Fig. 3F, G), in FRDA mice (bicep=92.46mA; FCR=70.60mA), compared to control animals (bicep 34.12mA; FCR=34.55mA). Additionally, at threshold EMG responses, FRDA mice produced substantially smaller MEP responses in both bicep and FCR muscles compared to age-matched controls (Fig. 3E). To enable a comparison of MEP responses between animal groups, recruitment curves were used to examine EMG responses following progressively larger evoking M1 stimuli (Fig. 3H,I). As expected, in control mice, the response MEPs increased in size in response to increasing M1 stimulation between threshold and 1.3 times the threshold (Fig. 3H,I). Importantly, the recruitment relation in FRDA animals was significantly decreased in both the biceps (Fig. 3H *p<0.05 at 1.3 times) and FCR muscles (Fig. 3I *p<0.05 at 1.2 and 1.3 times). Taken together, these data demonstrate a significantly weaker motor cortex-to-muscle output in FRDA animals when compared to controls.

### Anatomical changes in CSNs and the CST

Our results indicate a significant impairment in CSN/CST function as measured through behavioral and physiological output. This is consistent with the idea that FRDA also leads to pathological changes within CSNs. Alternatively, the reduction in EMG responses may stem from problems with the CST connections to spinal interneuons, or spinal motor neurons (MNs). To begin to examine these possibilities we performed a detailed anatomical assessment of the CSNs, the CST, and spinal cord and spinal MNs in two and three month old FRDA mice.

CSNs are located in lamina Vb of the motor cortex (Steward et al., 2021). As a proxy for overall motor cortex organization, we first measured the thickness of all cortical layers. We evaluated this by DAPI staining in conjunction with two lamina-specific markers, Cux1 (for layers I-Ⅳ) and CTIP2 (for layers Ⅴ- Ⅵ) (Nieto et al., 2004; McKenna et al., 2011). Given that FRDA mice have smaller brains than control mice (Fig. 3A-B), we normalized our measurements by expressing lamina width as a percentage of the total thickness of all cortical layers.

Total cortical thickness was reduced by 20 %(n=3) in FRDA mice. Layer I showed an increase in width, whereas layers Ⅱ/III and Ⅳ were significantly reduced in FRDA mice, yet there was no change in the width of layer V. This indicates that despite other cortical changes, there is no major organizational disruption in the CSN lamina (Fig. 4C-D). To Assess the number of CSNs in M1, we injected an adeno associated virus that expresses the tdTomato fluorescent protein (AAV-tdTomato) into the cervical spinal cord to retrogradely CSNs through the CST. No difference was observed in the number of back-labeled CSNs in FRDA mice (Fig. 4E-F). We did, however, observe a significant reduction in the area occupied by PKCγ^+^ CST axons at C6-8 level of the spinal cord in FRDA mice (Fig. 4G-H).

**Fig. 4.**
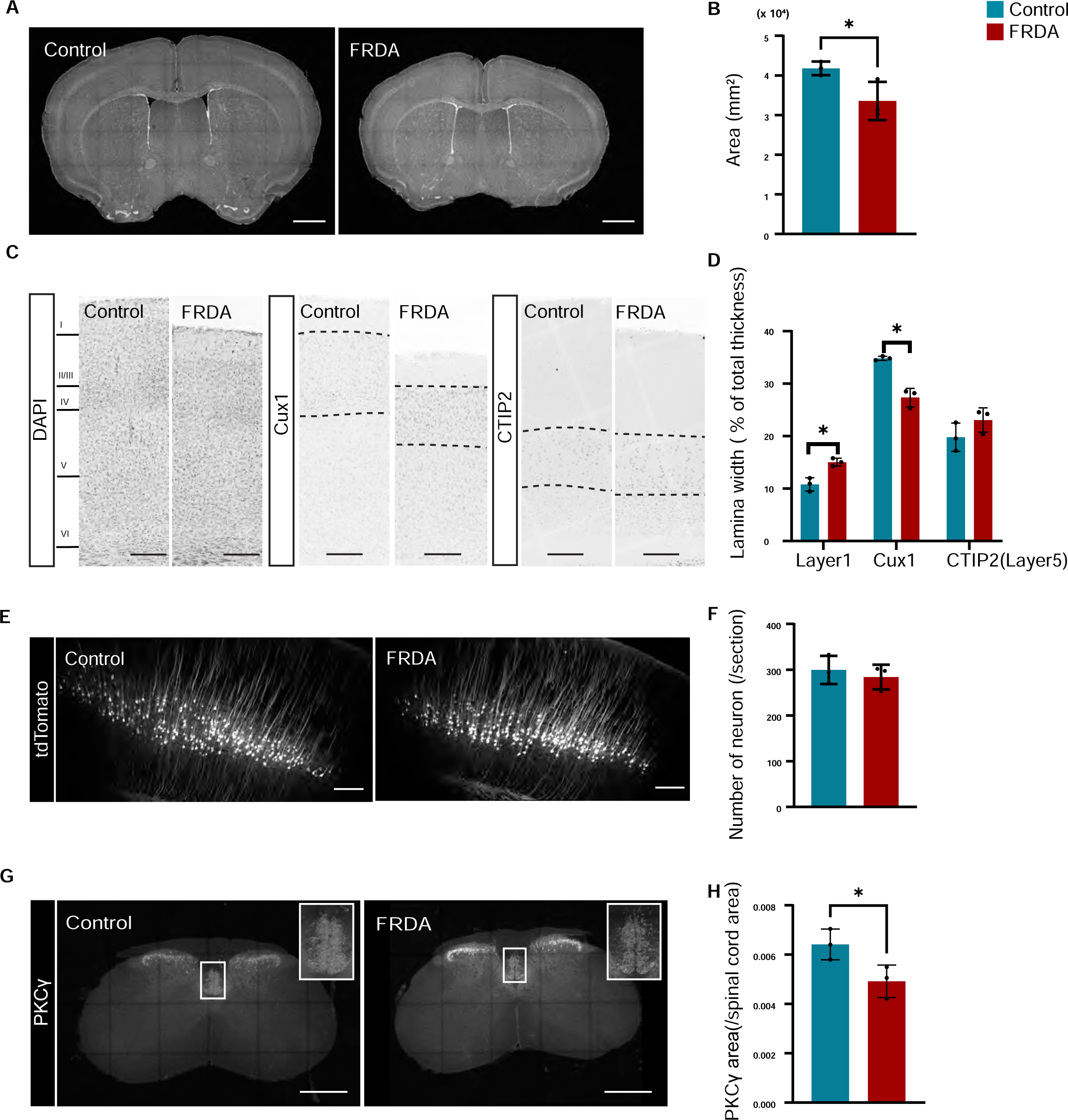
FRDA mice show anatomical changes in the CST. (A), DAPI staining in the brain in 2 months old Control (left) and FRDA (right) mice. (B), Brain area at 2 months old (n=3). C, Immunostaining of DAPI, Cux1, and CTIP2 in motor cortex. (D), Width of cortical layers (n=3). (E), tdTomato labeled corticospinal neurons in motor cortex. (F), Number of corticospinal neurons/section (n=3 mice). G, Immunostaining of PKCγ in C6-8 revel spinal cord. (H), Area of PKCγ positive CST in C6-8 level spinal cord (n=3). *p<0.05 (Student’s t test). All graphs are represented as Mean ± SD. Scale bars: 1 mm (A), 200 μm (C, E), 500µm (G).

We next examined the connectivity of CST axons at forelimb spinal cord. Consistent with the overall smaller size of FRDA mice, we also find that the spinal cord is reduced in size compared to control animals. When comparing cross sections of the spinal cord at C6-C8 between FRDA and control mice (Fig. 5A-B) we find that the ratio of the white matter to the to the overall spinal area was significantly reduced in FRDA mice, while the ratio of the gray matter to the total cross-section size was increased. We also assessed the number and size of spinal MNs using ChAT immunostaining Fig. 5C). However, we were unable to detect any differences in MN number or size across comparable C8 spinal sections. (Fig. 5D-E). Lastly, since CSNs are glutaminergic neurons, we evaluated the total number of vGluT1^+^ boutons in the spinal gray matter (Fig. 5F). Although many sensory neurons similarly rely on vGluT1 for their central transmission, we reasoned that a reduction in vGluT1^+^ CST boutons could still be detected this way. We normalized our measurements by measuring the number of vGluT1^+^ boutons per mm^2^ of gray matter. Nevertheless, when we performed these analyses, we observed no significant differences in the number of vGluT1^+^ boutons between FRDA and control animals (Fig. 5G). Thus, with exception of the thinning CST axons, FRDA mice show minimal abnormalities in the organization of the CSN/CST/spinal cord pathway.

**Fig. 5.**
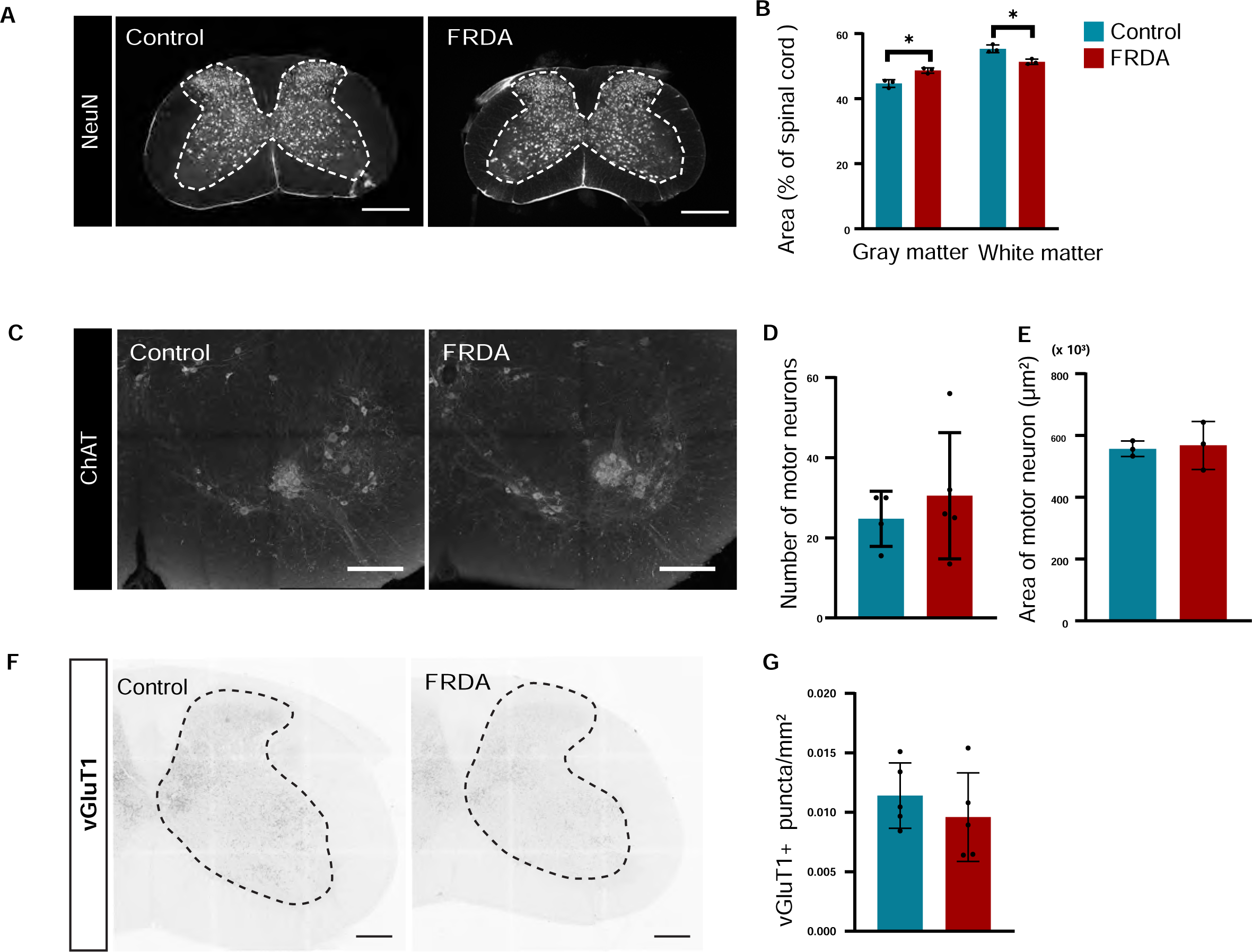
Anatomical changes in the spinal cord of FRDA mice. (A), Immunostaining of NeuN at C6-8 level spinal cord. (B), Area of gray matter, white matter, and total spinal cord (n=3). (C), Immunostaining of ChAT. (D), Number of ChAT positive motor neurons in each section. (E), Size (diameter) of ChAT positive motor neurons (n=5). (F), vGluT1 immunostaining in spinal cord. (G), Number of vGlut1^+^ puncta in each section at C8 level spinal cord (n=5).

### Changes in mitochondrial distribution in FRDA CSN neurons and CST axons

To investigate whether there are any alterations in the numbers or morphology of mitochondria in CSNs of FRDA mice, we selectively labeled the mitochondria of CSNs using AAVs (Fig. 6A). While there was no difference in the overall volume of mitochondria between control and FRDA mice, when corrected for cell body volume, CSNs in FRDA mice exhibited a higher proportion of mitochondria compared to CSNs in control mice (Fig. 6B-D). In addition, some FRDA mice showed large GFP-negative vacuoles in cytosol, nuclei, and/or dendrites (Fig. 6E). While the difference in the percentage of neurons with vacuoles between control and FRDA mice did not reach significance (Fig. 6F), we find that the vacuoles in CSNs in FRDA mice were larger than those in controls (Fig. 6G). FRDA mice also showed increased axonal swellings (Fig. 6H). Finally, when we assessed the distribution of the labeled CST associated mitochondria in spinal cord, we noted that the CST mitochondria are more abundant in spinal varicosities in FRDA mice when compared to controls (Fig. 6I, J). Together, these data indicate the presence of various morphological abnormalities in FRDA mice.

**Fig. 6.**
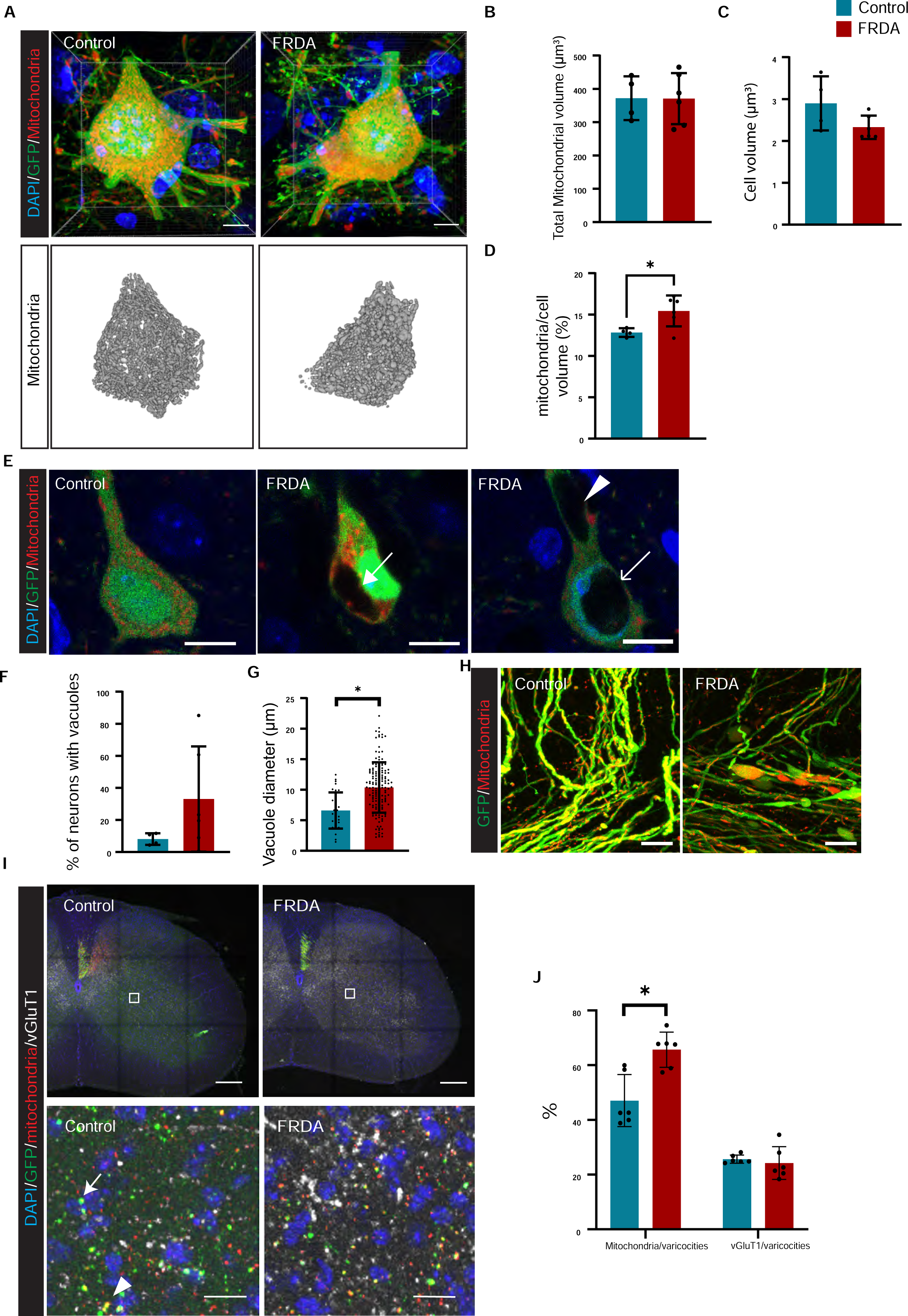
FRDA mice show vacuoles and altered mitochondrial distributions in CSNs (A), Labeling of DAPI (Blue), GFP (Green), and mitochondria (Red) in CSNs. (B-D), Bar graphs represents average of total mitochondrial volume in each neuron (B, n=4-6), average of cell body volume (C, n=4-6) and average of mitochondria/cell body ratio (D, n=4-6). (E, F), Representative images of GFP-negative vacuoles in FRDA mice. Arrow in E indicates GFP negative cytoplasmic vacuole. GFP-negative vacuoles are observed in cytosol (arrow in E center) dendrites (arrowhead in Eleft) and nucleus (arrow in Eleft). (F), Percentage of neurons that have GFP-negative vacuoles (n=4-6). (G), Diameter of GFP-negative vacuoles (n=4-6). (H), Representative images of axonal swellings in CSNs/CST. (I) Labeling varicosities of corticospinal axons using GFP (green), mitochondria (red), and vGluT1^+^ presynaptic boutons (white) in C6-8 spinal cord. (J), Bar graphs represent ratio of mitochondoria(arrow) and vGlut1(arrowhead) positive varicosities. *p<0.05 (Student’s t test, D, G, K). All graphs are represented as Mean ± SD. Scale bars: 5 µm (A), 10 µm (E,H), 200 µm (I upper pics), 15 µm (I lower pics).

## Discussion

FRDA causes progressive degeneration in the nervous system, most notably in DRG, Cerebellum, and CSNs/CST. Naturally, understanding when and how a loss of FXN affects these neural tissues is critical in the design and evaluation of therapeutic interventions. Yet, while pathological changes in DRG and Cerebellum have been fairly well characterized in FRDA, the CSNs/CST remain surprisingly understudied. We here investigated the impact of Fxn loss on CSNs/CST function and morphology in the recently described YG8JR FRDA mouse model.

Several lines of evidence, including data from this study, support the idea that the YG8JR mouse model represents one of the better preclinical models of FRDA. The median age of onset of disease in patients is 10.52 years old (Harding, 1981) and as shown previously, YG8sR mice developed motor dysfunction at 2-3 months old (Gérard et al., 2023; Kalef-Ezra et al., 2023). Similarly, our studies detected significant motor deficits starting as early as two months of age, which in humans corresponds to a juvenile stage (Wang et al., 2020). It is possible that the motor deficit may be present even earlier, but assessing this at earlier ages is less reliable, especially when there are such vast differences in weight between FRDA and control animals. The observed weight loss in FRDA mice may be an indirect consequence of a reduced motor capability, however, as the motor deficits may disadvantage mutant pups when competing for nursing time. (A condition which naturally does not apply to FRDA newborn babies). Thus, further studies may be needed to better assess the time of onset of motor deficits in YG8JRex4 FRDA mice.

Another possible similarity between the YG8JRex4 mouse model and FRDA patients are the gastro-intestinal abnormalities we observed. By seven months of age, 70% of male mice exhibited rectal prolapse. While intestinal impairments are not among the most debilitating symptoms in FRDA, clinical reports have shown that the prevalence of diseases such as Crohn’s disease, ulcerative colitis, and irritable bowel syndrome is 2 to 3.6 times higher in FRDA patients compared to the non-FRDA population (Shinnick et al., 2016). In addition, it was reported that 64% of patients complain of bowel symptoms (Lad et al., 2017). The immediate cause of these problems remains poorly understood but could relate to a loss of sensory innervation of the intestinal wall (Servin-Vences et al., 2023; Wolfson et al., 2023) or to body wall muscle weakness (Yiou et al., 2001). The rectal prolapse phenotype was occasionally also observed in a different laboratory (J. de Nooij, personal communication) but was not reported in previous other studies of the YG8JR mice. This could indicate that this phenotype may be sensitive to environmental factors. Alternatively, this could reflect a difference between the mouse *Fxn* exon 2 (prior studies) and exon 4 (this study) deletion mutants. Nevertheless, based on our observations, the YG8JRex4 mouse model may offer an opportunity to better understand the role of Fxn in gastrointestinal function.

Neuronal dysfunction of large caliber proprioceptive DRG sensory neurons is one of the first hall marks in FRDA and can be readily recognized through the loss of deep tendon reflexes (Geoffroy et al., 1976). Using a battery of motor behavioral test, we here show a general reduction in locomotor ability, with FRDA animals exhibiting reductions is locomotor distance, speed, as well as grip strength. Unlike prior studies, we did not observe differences in Rotarod test between control and FRDA mice. It is also important to note that the smaller body size and weight of FRDA mice could potentially influence the results of these behavioral tests. Besides general motor deficits, FRDA animals also showed significant deficits in digit control, which is essential for skilled motor behaviors. For instance, FRDA mice were severely impaired in grabbing onto a food pellet or manipulating a small pasta stick. These tests are equivalent to the nine-peg and block test, which are commonly performed with FRDA patients to assess fine motor skill. Damage to the CST results in poorer scores on pellet reaching and pasta handling tests (Khaing et al. 2012). Therefore, the poor performance of FRDA mice on these motor tasks is consistent with a possible functional impairment of the CSNs or the CST. Indeed, we observed a profound functional loss in CSN output when we assessed motor endplate potentials in forelimb muscle following motor cortex stimulation. Nevertheless, we cannot exclude that the observed motor phenotypes are solely due to CSN/CST dysfunction since impaired proprioceptive or cerebellar function may have similar consequences. To investigate the primacy of CSNs and the CST in the FRDA related loss of skilled motor function it will be required to selectively eliminate Fxn from CSNs.

Our morphological studies to determine the cause of CSN/CST dysfunction in FRDA animals indicated normal numbers of CSNs at three months of age, however, we observed several potential causes of CSN/CST dysfunction. First, we noted a decrease in the area normally occupied by PKCγ^+^ CST axons at spinal segmental levels. This could indicate thinner CST axons or it could indicate a reduction in myelination of CST axons such that they are more compact. Second, we observed large mitochondria-free vacuoles in CSN cell bodies in FRDA mice (Fig. 6). These vacuoles resemble those reported in DRG of other FRDA mouse models (Anjomani Virmouni et al. 2015; Kemp et al. 2017; Sandi et al. 2011; Tomassini et al. 2012; Al-Mahdawi et al. 2006). Similar to previous reports, vacuoles were observed in both the cytoplasm and the nucleus. Additionally, we detected vacuoles in dendrites. Previous reports using electron microscopy analyses have speculated that the vacuoles observed in the DRG of FRDA mice are autophagosomes (Simon et al. 2004). To confirm whether CSN vacuoles similarly may constitute autophagosomes, EM studies or an immunological analysis with autophagosomal markers will be required. The nuclear vacuoles remain to be characterized but are unlikely to be autophagosomes. Interestingly, however, in adenosine deaminase acting on RNA 2 (ADR2) knockout mice, a model of amyotrophic lateral sclerosis (ALS), motor neurons also show nuclear vacuoles (Sasaki et al. 2014). In these mice it has been suggested that this is caused by a calcium imbalance, given that the loss of ADR2 increases the proportion of calcium-permeable AMPA receptors. Previous studies have implicated Fxn not just in iron homeostasis but also in regulating Ca^2+^ homeostasis (Tamarit et al. 2021). Thus, possibly the nuclear vacuoles we observed in CSNs may be a consequence of a calcium imbalance due to Fxn loss.

The third morphological defect we noted in in FRDA mutant mice were the CST axonal swellings. Such axonal spheroids or swellings were similarly reported in Purkinje cells in FRDA patients and in DRG sensory neurons in FRDA mice (Kemp et al. 2016; Mollá et al. 2017). Axonal swellings may be a general sign of neuronal degradation given that they have also been observed in traumatic brain injury and in neurodegenerative diseases such as Alzheimer’s disease or Multiple Sclerosis (Reeves, Phillips, and Povlishock 2005; Nikić et al. 2011; Krstic and Knuesel 2013). The axonal swellings in CSNs/CST were packed with mitochondria. This finding is also consistent with observations in cultured primary sensory neurons isolated from FRDA mice. The abnormal axonal swellings could potentially influence the proper transmission of action potentials (Wu, Gilpin, and Adnan 2020). If so, this would align with our electromyography (EMG) observations, which demonstrated that stronger stimuli were required to elicit muscle responses in FRDA mice (Fig. 3).

Taken together, our studies show significant CSN/CST dysfunction which appears to be routed in various morphological aberrations that may be indicative of neuronal degeneration, including the thinning of CSN axons, the increased incidence of cytoplasmic and nuclear vacuoles, and axonal swellings. These changes in CSNs resembles observations in DRG and cerebellum in FRDA mice and patients, which could suggest that Fxn loss may result in similar cellular defects in all three tissues. Moreover, we find that these abnormalities can be observed as early as 8-12 weeks of mouse age, indicating that FRDA-associated CSN/CST impairments may arise earlier in the disease process than previously thought. As such, these findings offer valuable new insights in the pathophysiology of FRDA, and consequently, for the development and evaluation of therapeutic strategies.

## Acknowledgement

This work was supported by the Structural and Functional Imaging Core at Burke Neurological Institute and S10 Shared Instrumentation Grant OD028547-01. Y.Y. was supported by National Institute of Neurological Disorders and Stroke Grants NS100772, NS115963, NS119508, and NS093002, and Friedreich’s Ataxia Research Alliance. M,N was supported by Friedreich’s Ataxia Research Alliance, The Uehara Memorial Foundation, and Mochida Memorial Foundation for Medical and Pharmaceutical Research. We thank J. Magrane (Will Cornell Medicine) for providing *Fxn*^Δ4/+^ mice. We thank I. Pavlova (Burke Neurological Institute) for technical advice of confocal images; W. Duggan and A. Bernstein (Burke Neurological Institute) for making chamber for pasta handling test. We thank S. Cho and I. Kim (Burke Neurological Institute) for letting us use open field apparatus.

## Notes

### Competing Interest Statement

The authors have declared no competing interest.

